# Scale drop disease virus (SDDV) triggering ferroptosis both *in vivo* and *in vitro* facilitates virus infection via targeting transferrin receptor 1 (TfR1)

**DOI:** 10.1101/2025.03.18.643978

**Authors:** Jiaming Chen, Yuting Fu, Shuyi Chen, Weng Shaoping, Jianguo He, Chuanfu Dong

**Affiliations:** State Key Laboratory of Biocontrol/School of Life Sciences of Sun Yat-sen University, Guangzhou 510275, China; Southern Marine Science and Engineering Guangdong Laboratory (Zhuhai), Zhuhai 519000, China; Zhuhai Modern Agriculture Development Center, Zhuhai 519000, PR China; Institute of Aquatic Economic Animals and Guangdong Province Key Laboratory of Aquatic Economic Animals, Sun Yat-sen University, Guangzhou 510275, People’s Republic of China

**Author notes:** Corresponding author, Chuanfu Dong, Ph.D, School of Life Sciences, Sun Yat-sen University, No. 135, Xingang Road West, Guangzhou, 510275, P. R. China., E-mail addresses.

**Keywords:** Scale drop disease virus (SDDV), Ferroptosis, Transferrin receptor protein 1, Infectivity

## Abstract

Scale drop disease virus (SDDV) is a distinct member in genus *Megalocytivirus* of family *Iridoviridae*, garnering increasing attention due to its significant threat to teleost. Ferroptosis is a new type of cell death discovered recently and involved in various viral infections. Knowledges on SDDV induced ferroptosis remains unclear. Here, we demonstrated that SDDV infection triggers ferroptosis, as evidenced by hallmark features such as iron overload, massive lipid peroxides accumulation, glutathione depletion and glutathione peroxidase 4 downregulation. SDDV-infected MFF-1 cells exhibited increased reactive oxygen species production and mitochondrial shrinkage. Treatment with Ferrostatin-1, a potent ferroptosis inhibitor, significantly attenuated SDDV replication in MFF-1 cells and could improve the survival of mandarin fish upon SDDV challenge. Treatment with an iron chelator mitigated ferroptosis and reduced the mortality of mandarin fish following SDDV infection, suggesting that SDDV-induced ferroptosis is iron-dependent. Finally, we demonstrated that SDDV infection could upregulate the expression of transferrin receptor protein 1 (TfR1), a critical iron transporter, to disrupt cellular iron homeostasis, induce ferroptosis, and then facilitate viral infection. Collectively, our findings provide compelling evidence that SDDV infection induces ferroptosis by targeting TfR1 to facilitate virus infection. Inhibiting ferroptosis represents a promising immunotherapeutic strategy for combating SDDV infection in aquaculture.

## 1. Introduction

Scale drop disease virus (SDDV) is a sense double-stranded large DNA virus belonging to the *Megalocytivirus lates1* species of genus *Megalocytivirus* within subfamily *Alphairidoviridae*. Since it was first scientifically identified in 2015 [1], SDDV has been widely reported in Asian seabass *Lates calcarifer* industry across several Southeast Asian countries, including Singapore, Malaysia, and Thailand, and has emerged as one of the most concerning viral pathogens to farmed Asian seabass in these regions [2-4]. SDDV can infect Asian seabass of various sizes, especially threatening large-sized ones, with accumulative mortalities from 40% to 50% [1, 4]. The documentation of SDDV-associated diseases, known as scale drop syndrome (SDS), could date back to as early as in the early-1990s in Asian seabass farms in Penang, Malaysia [4, 5]. Different from SDDV in SE countries just infecting Asian seabass, SDDV was isolated from yellowfin seabream *Acanthopagrus latus* suffering from ascites diseases in mainland China by our team in 2020 [6]. Furthermore, our recent study showed that even a low dose of SDDV could cause 100% mortality in mandarin fish *Siniperca chuatsi*, a highly susceptible host fish species to broad *Megalocytivirus* infections [7, 8], suggesting that mandarin fish may serve as a promising infection and vaccination model for the study of SDDV [9]. Additionally, SDDV-close viruses, including European chub iridovirus (ECIV) and a novel tilapia megalocytivirus, have also been reported in European chub *Squalius cephalus* in UK [10] and tilapia *Oreochromis* spp. in the USA [11], respectively. Collectively, all these findings highlight the significant threat of SDDV to global aquaculture and underscore the need for heightened vigilance.

Besides SDDV, the genus *Megalocytiviruses* also includes another species, namely *Megalocytivirus spagrus1*, which is represented by the type isolate of infectious spleen and kidney necrosis virus (ISKNV) [12]. Nowadays, *Megalocytivirus spagrus1* is further divided into three genotypes: ISKNV, red sea bream iridovirus (RSIV) and turbot reddish body iridovirus (TRBIV). These viruses have caused lethal systemic infections in wild and farmed freshwater, brackish, and marine fish species worldwide [6, 7, 13]. Compared to the extensively studied ISKNV, studies on SDDV remain at a relatively preliminary stage, with the mechanisms underlying its pathogenesis still largely unexplored.

Ferroptosis is a novel nonapoptotic form of programmed cell death and caused by lipid peroxidation (LPO) due to the accumulation of iron-dependent reactive oxygen species (ROS) [14]. It is characterized by the deactivation of the cellular antioxidant system, primarily through the depletion of glutathione (GSH) and the functional loss of phospholipid peroxidase glutathione peroxidase 4 (GPx4) [15]. Notably, these hallmarks of ferroptosis have been observed during various viral infections, suggesting a close association between ferroptosis and viral pathogenesis. For instance, ferroptosis is linked to severe acute respiratory syndrome coronavirus 2 (SARS-CoV-2) infection and may lead to multi-organ damage [16]; Newcastle disease virus (NDV) has been shown to induce ferroptosis to kill tumor cells [17]; and Herpes Simplex Virus 1 (HSV-1)-induced ferroptosis contributes to viral encephalitis [18]. Additionally, a variety of viruses, including Influenza A Virus (IAV) [19], Rotavirus [20] and Porcine Reproductive and Respiratory Syndrome Virus (PRRSV) have been reported to trigger ferroptosis to enhance viral replication [21].

Transferrin receptor 1 (TfR1) is a type II transmembrane glycoprotein consisted of a disulfide-bonded homodimer and functions as the primary cellular iron absorption protein. TfR1 binds transferrin (TF) at the cell surface, importing iron from the extracellular environment into cells via receptor-mediated endocytosis [16]. Imbalance in iron metabolism is regarded as a crucial factor in ferroptotic cell death. Thus, TfR1 is considered to be an important regulator of ferroptosis [22]. Previous studies have shown that TfR1 plays a key role in ferroptosis induced by viral infections. For example, upregulation of TfR1 promotes ferroptosis during coxsackievirus B3 (CVB3) infection [23]. And dysregulation of TfR1 and TF contributes to swine influenza virus (SIV)-induced ferroptosis, enhancing viral replication [24]. However, research on the role of ferroptosis in aquatic viral infections remains limited, with little known about whether aquatic viruses mediate ferroptosis or how ferroptosis influences viral replication in aquaculture species.

In this study, we determined that SDDV infection induces ferroptosis and TfR1 is identified as a key regulator of iron overload during SDDV infection, leading to ferroptotic cell death. These findings provide a novel perspective on SDDV pathogenesis and offer theoretical support for the development of potential antiviral therapeutic strategies to mitigate SDDV infection.

## 2. Materials and methods

### 2.1. Cell culture and viral infection

Mandarin fish fry (MFF-1) cell line was established in our laboratory and used for virus isolation and infection studies [25]. MFF-1 cells were grown in complete Dulbecco’s Modified Eagle’s Medium containing 10% fetal bovine serum (FBS), penicillin (100 IU/ml), streptomycin (100 μg/ml), and amphotericin B (0.25 μg/ml; Life Sciences, USA) at 27 °C.

SDDV ZH-06/20 strain was isolated by our team and stored at −80 °C [6]. MFF-1 cells were incubated with ZH-06/20 at a multiplicity of infection (MOI) of 5.0 for 1 h in serum-free DMEM. Then, the cells were washed with phosphate-buffered saline (PBS) and replenished with fresh DMEM containing 5% FBS. Uninfected MFF-1 cells served as the mock control group. Cells were harvested at 6, 12, 24, 36, 48 hours post infection (hpi) for subsequent experiments. All experiments were conducted in triplicate.

### 2.2. Artificial infection and experimental groups

Mandarin fish (50±10 g) were obtained from aquaculture farmers in Nanhai District, Foshan, Guangdong Province, China. The fish were temporarily reared in a 500 L plastic tank. During the experiment, individuals were randomly assigned to experimental groups. Fish in the SDDV group were intraperitoneally injected with diluted SDDV ZH-06/20 solution dissolved in normal saline (10^3^ TCID_50_/fish), whereas control group fish received an equivalent volume of normal saline. Tissue samples were collected at 2, 4, 6, 8, 10, 12, 14 days post-infection (dpi) from each group for morphological, biochemical and molecular analyses. This study was approved by the Institutional Animal Care and Use Ethics Committee of Sun Yat-sen University under number 0017097003.

### 2.3. Quantitative real time PCR (qRT-PCR)

Primers for qRT-PCR were designed using NCBI Primer-BLAST (Table 1). The *β-actin* gene was used as the internal reference for normalization. Total RNA was extracted and reverse-transcribed into complementary DNA (cDNA) using Evo M-MLV RT Premix for qPCR (Accurate Biology, Hunan, China). The qRT-PCR assays were performed on a LightCycler® 480-II Multiwell Plate 384 real-time detection system (Roche Diagnostics, USA). The qRT-PCR reaction volume included 10 μL of SYBR qPCR Master Mix (Accurate Biology, Hunan, China), 2 μL of cDNA, 0.5 μL of each primer pair, and 7 μL ddH_2_O. The amplification reaction was conducted at 95 °C for 1 min, 1 cycle; 40 cycles of denaturation at 95 °C for 10 s and annealing at 60 °C for 30 s; 95 °C for 5 s, 60 °C for 1 min and 95 °C, 1 cycle; 50 °C for 30s, 1 cycle. All experiments were conducted in triplicate. Relative expression levels were analyzed using the 2^-ΔΔCt^ method.

**Table 1.**
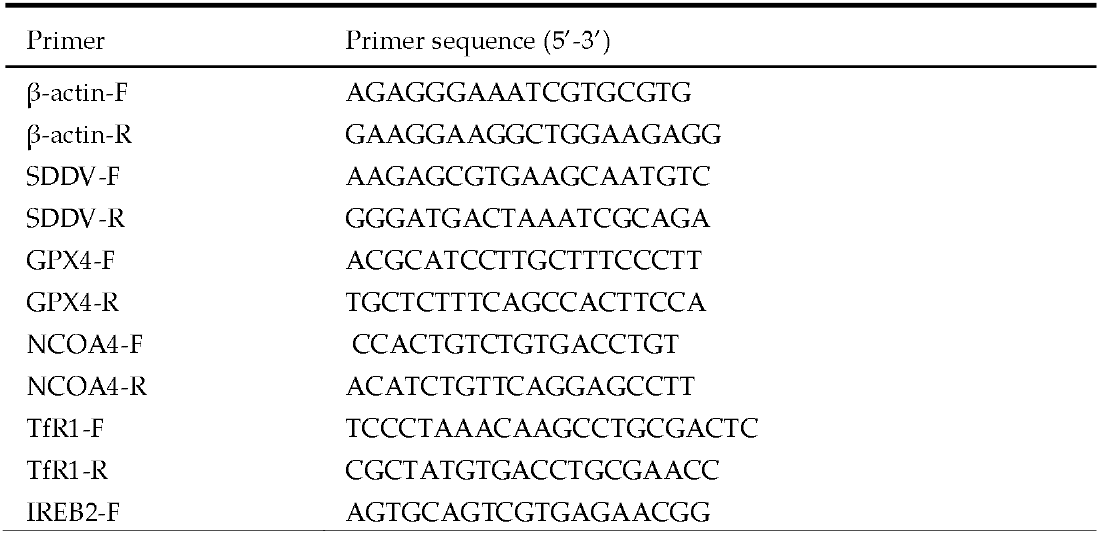
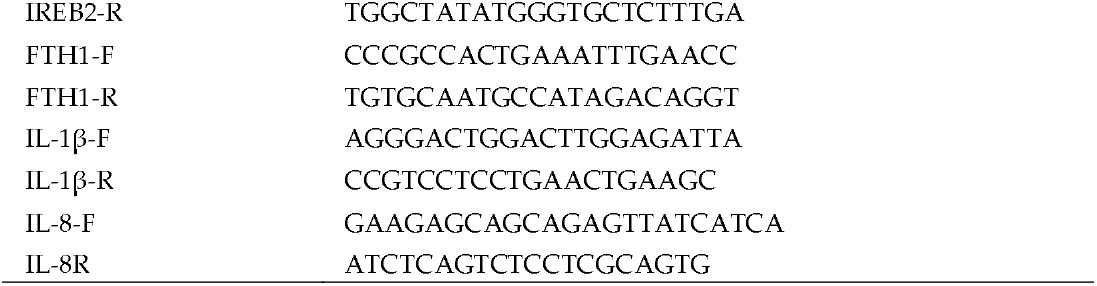
Primers used in this study

### 2.4. Immunofluorescence assay (IFA)

An IFA was performed to confirm SDDV infection. A monoclonal antibody (mAb) against SDDV major capsid protein (MCP) (1:2,000) was used as the primary antibody. Alexa Fluor 555-conjugated donkey anti-mouse IgG (Invitrogen, Carlsbad, California, USA) was used as the secondary antibody. SDDV-infected MFF-1 cells served as the positive control, while PBS-treated cells were used as the negative control. Nucleus were stained with DAPI (Abcam, Shanghai, China), and sections were observed under a fluorescence microscope microscopy.

### 2.5. Prussian blue staining section observation

Spleen tissues from mandarinfish infected with the ZH-06/20 strain were collected and fixed in alcohol-formalin-acetic (AFA) solution before embedding in paraffin. Embedded tissues were excised into thickness of 3 μm, dewaxed in xylene, rehydrated in graded ethanol, and stained with Prussian blue for 10–20 minutes following standard protocols [26]. Observations were made under a microscope (Nikon, Tokyo, Japan).

### 2.6. Transmission electron microscopy (TEM)

TEM analysis of SDDV-infected MFF-1 cells was performed at 60 hpi, as described previously [23]. Briefly, infected cells were fixed in 2.5% glutaraldehyde and 0.1 M PBS containing 2.0% osmium tetroxide. After dehydrating, penetrating, embedding and polymerization, the samples were cut into ultrathin sections of 60 nm. Sections were stained with uranyl acetate-lead citrate and examined under a Philips CM10 electron microscopy (FEI Company, Hillsboro, Oregon, USA).

### 2.7. Iron assay

The total iron concentration was determined using an iron assay kit (Solarbio, Beijing, China) in accordance with the manufacturer’s instructions. Briefly, approximately 0.1 g of tissue was homogenized in 1 mL of extraction buffer on ice. In MFF-1 cells, after removing the supernatant, the cells were digested with trypsin and collected. Subsequently, 1 mL of extraction buffer was added, and the cells were subjected to sonication on ice for disruption. The homogenate was centrifuged at 4,000 × g for 10 min at 4 °C, and the supernatant was collected. For the measurement tube, 60 μL of reagent 1, 120 μL of reagent 2, and 120 μL of the sample were added. A blank tube was prepared with 120 μL of distilled water, and a standard tube was prepared with 120 μL of gradient-diluted standard solution. Subsequently, 60 μL of chloroform was added to each tube, followed by incubation in a boiling water bath for 5 min. Afterward, the samples were centrifuged at 10,000 × g for 10 min. The optical density (*OD*) of each sample was measured at a wavelength of 520 nm. The total iron concentration was calculated based on the provided formula.

### 2.8. Fe2+ content assay

The Fe^2+^ concentration in tissues or cells was measured using a Fe^2+^ concentration assay kit according to the manufacturer’s instructions. Approximately 0.1 g of tissue was weighed and 1 mL of reagent 1 was added. The tissue was homogenized and then centrifuged at 10,000 × g for 10 min at 4°C. For MFF-1 cells, the supernatant was removed, and the cells were digested with trypsin, followed by collection of the cells and sonication on ice. 10 μL of a 40 mmol/L standard solution was added to 990 μL of distilled water and mixed thoroughly to prepare a 400 μmol/L standard solution. The standard solution was then serially diluted with reagent 1 to obtain gradient concentrations, which were prepared fresh for each experiment. The blank tube received 200 μL of ddH_2_O, the standard tubes were added with 200 μL of the appropriately diluted standard solutions, and the sample tubes received 200 μL of the prepared samples. Then, 100 μL of reagent 2 was added to each tube and incubated at 37 °C for 10 min. Afterward, 100 μL of chloroform was added to each tube. The tubes were vortexed for 5 min and centrifuged at 12,000 × g for 10 min. The absorbance at 593 nm was measured using a spectrophotometer. A standard curve was plotted, and the Fe^2+^ concentration was calculated using the corresponding formula.

### 2.9. Glutathione (GSH) assay

Intracellular GSH levels were determined using the GSH assay kit (Beyotime, Shanghai, China) following the manufacturer’s protocol. Spleen tissues were frozen in liquid nitrogen and ground into into powder. The cell sample was digested with trypsin and then centrifuged to remove the supernatant. Then, 30 μL of removal reagent M solution and 70 μL of reagent M solution were added, and the mixture was thoroughly homogenized. The homogenate was centrifuged at 10,000 × g for 10 min at 4 °C, and the supernatant was used for total GSH measurement. Absorbance was measured at 412 nm using a microplate reader (BIO-RAD Instruments, California, USA).

### 2.10. Lipid peroxidation (LPO) assay

The LPO content in tissues was determined using an LPO content assay kit (Solarbio, Beijing, China) according to the manufacturer’s instructions. Briefly, 0.1 g of tissue was homogenized in 1 mL of extraction liquid, followed by centrifuged at 8,000 × g for 10 min. Reagents 1, 2, and 3 were added according to the protocol. For the blank tube, distilled water was used, and for the standard tube, gradient-diluted standard solution was prepared. The samples were incubated in a water bath at 100 °C for 60 min and centrifuged at 8000 × g for 10 min. Absorbance values were measured at 532 nm and 600 nm, and LPO content was calculated using the provided formula.

### 2.11. 8-hydroxy-2’-deoxyguanosine (8-OHdG) assay

The concentration of 8-hydroxy-2’-deoxyguanosine (8-OHdG) in tissues was measured using an 8-OHdG assay kit (Jonlnbio, Shanghai, China) in accordance with the manufacturer’s guidelines. This kit utilizes a competitive enzyme-linked immunosorbent assay (ELISA). The 8-OHdG antibody was pre-coated onto a microplate. Samples or standards, and HRP-labeled antigens were sequentially added, followed by incubation and washing steps. The colorimetric substrate TMB was added, transitioning from blue to yellow upon acidification. The color intensity, inversely proportional to the 8-OHdG concentration, was measured at 450 nm using a microplate reader. Sample concentrations were calculated from the standard curve.

### 2.12. Reactive oxygen species (ROS) assay

ROS levels in MFF-1 cells were detected using a ROS assay kit (Beyotime, Shanghai, China), with the fluorescent probe DCFH-DA. The probe was diluted 1:1000 and added to adherent MFF-1 cells, then incubated for 20 min. The GFP fluorescence intensity was observed under a fluorescence microscope.

### 2.13. Western blot analysis

Protein samples were extracted from MFF-1 cells using RIPA lysis buffer (Beyotime, Shanghai, China) and quantified with the Pierce BCA Protein Assay Kit (Thermo Fisher Scientific, Massachusetts, USA). Proteins were separated via 10% sodium dodecyl sulfate-polyacrylamide gel electrophoresis (SDS-PAGE) and transferred onto Immobilon-P PVDF membranes (Millipore, Merck, Germany). The anti-SDDV MCP mAb (1:2,000) or rabbit polyclonal antibodies (pAbs) against TfR1 (1:2,000) was used as primary antibody, with β-actin as the internal control. HRP-conjugated goat anti-rabbit or anti-mouse IgG was used as secondary antibody. The blot was visualized using the Tanon High-sig ECL Western Blotting Substrate (Tanon, Shanghai, China).

### 2.14. RNA interference (RNAi)

Small interfering RNAs (siRNAs), including a non-targeting control siRNA and siRNA targeting TfR1, were synthesized by RiboBio (Guangzhou, China). The siRNAs were transfected into MFF-1 cells using the jetPRIME transfection reagent (Ployplus, Strasbourg, France) following the manufacturer’s protocol. Briefly, 2.5 μL of siRNA was diluted in 50 μL of jetPRIME buffer and mixed with 1 μL of jetPRIME transfection reagent. The reaction mixture was incubated at room temperature for 10 min, and then added to the culture medium in 24-well plates. The final siRNA concentration was adjusted to 100 nM. Forty-eight hours post-transfection, TfR1 mRNA expression levels were assessed by qRT-PCR, and protein levels were evaluated by western blot analysis. After 48 hours post-transfection, cells were infected with the ZH-06/20 at a MOI of 5.0. The cells were collected at6, 12, 24, 36, 48, 60 and 72 hpi for subsequent analyses. The siRNA sequence targeting mandarinfish TfR1 was as follows: ACT CTC AAG GCT CTG ATC A.

### 2.15. Drug treatment

Ferrostatin-1 (Fer-1) was purchased from GLPBIO (California, USA) and erastin, ferric citrate (FAC), deferoxamine mesylate (DFO), necrostatin-1 (Nec-1) and Z-VAD were obtained from MedChemExpress (New Jersey, USA). In vitro, these compounds were diluted in dimethyl sulfoxide (DMSO) to prepare working solutions at the desired concentrations and added to the cell culture medium 5 h before SDDV infection. In vivo, the drugs were diluted with DMSO to appropriate concentrations and administered to mandarinfish via intramuscular injection 5 h prior to SDDV infection.

### 2.16. Cell viability assay

MFF-1 cells were seeded into 96-well plates at a density of 5 × 10^3^ cells/well and cultured for 24 h. The test drugs were prepared at various concentrations, added to the culture medium, and incubated with the cells for 5 h. Subsequently, Cell Counting Kit-8 (CCK-8) solution (Beyotime, Shanghai, China) was introduced into each well and incubated at 27 °C for 1 h. Cell viability was assessed by measuring the *OD* at 450 nm using a microplate reader. Viability data were normalized relative to the control group.

### 2.17. Statistical

The phenotypic characteristics of cells were assessed using a series of assays, with statistical analyses conducted as follows: Student’s *t*-test was employed to evaluate differences between groups, while Pearson’s correlation test was utilized to examine relationships between variables. Analysis of variance (ANOVA) was performed to compare multiple groups, focusing on both inter-group and intra-group variances. Notably, the inter-group variance ratio was relatively small. Statistical significance was defined as *p* < 0.05. All data analyses were conducted using ORIGIN 2021 software.

## 3. Results

### 3.1. SDDV but not ISKNV and MRV induces classical ferroptosis in mandarinfish

To explore whether SDDV induces ferroptosis *in vivo*, we applied the SDDV-mandarinfish model established by our team and examined the features of ferroptosis in spleen tissue, the primary target of SDDV infection [9]. Accumulation of cellular iron is one of the major biochemical hallmarks of ferroptosis. Thus, the tissue sections of infected mandarinfish were stained with Prussian blue to visually observe the changes of Fe^3+^. Additionally, the levels of Fe^2+^ and total iron were quantified using specific assay kits. ISKNV, a type species in *M. spagrus1* and MRV, a distinctive member of genus *Ranavirus* in the *Iridoviridae* family [27, 28], were used as controls. As shown in **Fig. 1A**, hemosiderin accumulation was observed in the spleen tissue of mandarinfish infected with SDDV, ISKNV, and MRV. The contents of total iron and Fe^2+^ increased significantly after SDDV and ISKNV infection compared to the control group, whereas no significant increase was observed in the MRV group (Fig. 1B). Ferroptosis is also characterized by increased lipid peroxidation (LPO) induced by reactive oxygen species (ROS) and the dysfunction of antioxidant defense mechanisms. Compared to the control group, LPO levels were significantly elevated (Fig. 1C), GSH levels were decreased, reaching the lowest at 12 dpi (Fig. 1C). GPx4 expression was also significantly down-regulated in the spleen tissue following SDDV infection (Fig. 1C). The levels of 8-hydroxy-2-deoxyguanosine (8-OHdG), a marker of oxidative DNA damage caused by ROS, were markedly elevated following ZH-06/20 infection (**Fig. 1C**). In contrast, although 8-OHdG levels increased in the ISKNV-infected group, LPO levels remained unchanged and even significantly decreased in the early stages of infection (**Fig. 1C**). Additionally, the content of GPX4 underwent a significant up-regulation at 3 and 6 dpi in the ISKNV infection group (**Fig. 1C**). After MRV infection, the levels of LPO increased significantly (Fig. 1C), while substantial changes were observed in total iron content (**Fig. 1B**), and 8-OHdG levels decreased following infection. Taken together, these results demonstrated that ferroptosis is present in SDDV-infection but is not apparent in ISKNV and MRV infections.

**Fig. 1.**
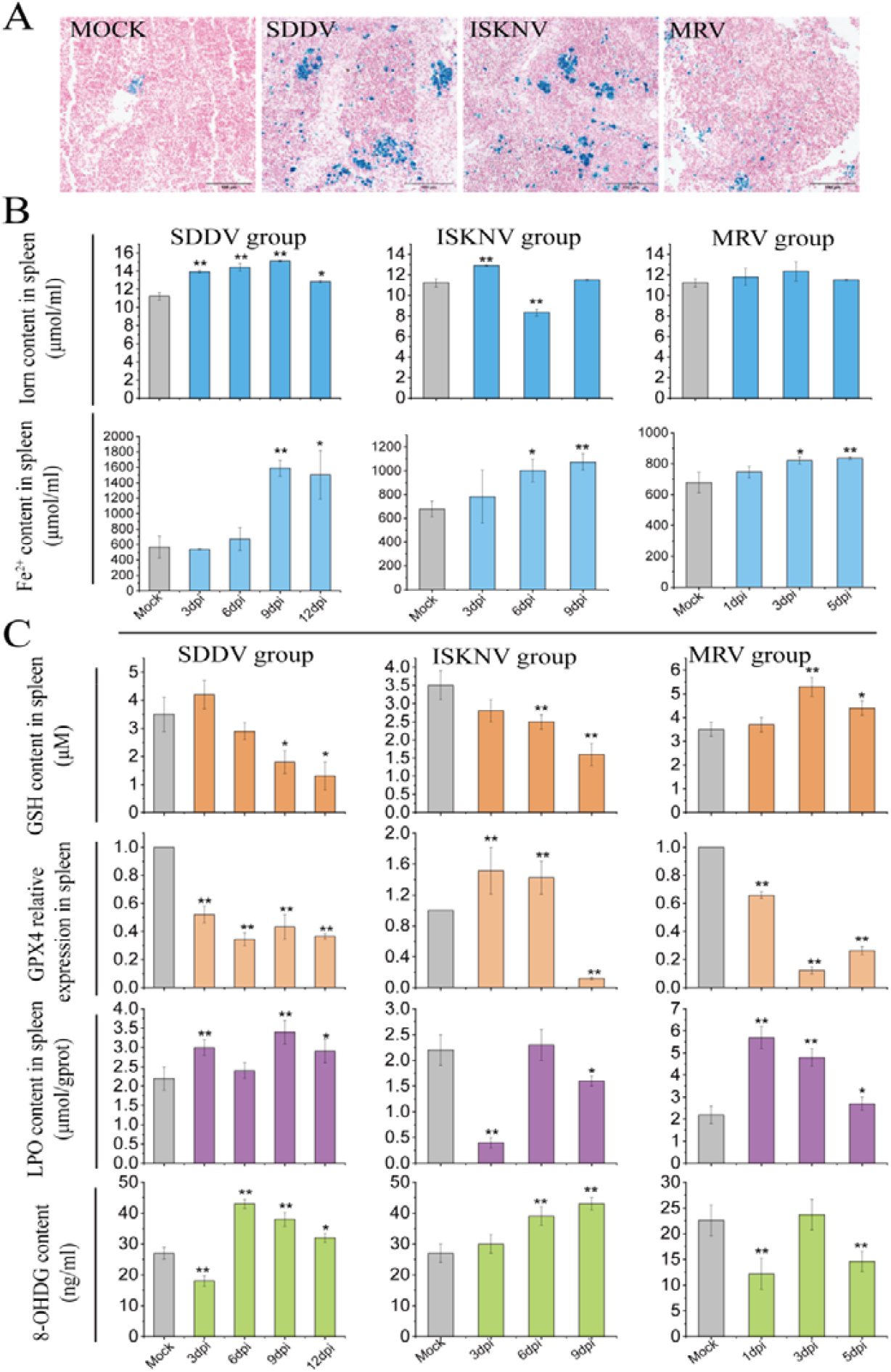
Ferroptosis occurs *in vivo* in mandarin fish infected with SDDV. **A**, Representative images of Prussian blue staining in spleen tissues from mock, SDDV, ISKNV, and MRV groups (scale bar = 50 μm). **B** Total iron and Fe^2+^ concentrations were measured in spleen tissues from mock, SDDV, ISKNV, and MRV groups using an iron assay kit. **C** The concentrations of GSH, LPO, and 8-OHdG were quantified using commercial assay kits following infections with SDDV, ISKNV, and MRV. The relative expression of GPX4 was analyzed by qRT-PCR. n = 3. *p < 0.05.

### 3.2. SDDV also triggers ferroptosis in MFF-1 cell

MFF-1 cell line is highly susceptible to SDDV infection and is widely used for the isolation, identification and infection mechanism research of iridoviruses [25]. To determine whether SDDV triggers ferroptosis in vitro, MFF-1 cells were infected with SDDV. Firstly, we evaluated the morphological changes of mitochondria in MFF-1 cells infected with SDDV at an MOI of 5.0 for 72 h, as mitochondrial shrinkage is a hallmark of ferroptosis [29]. Infected cells exhibited disrupted cristae and shrinkage of mitochondrial compared to uninfected cells **(Fig. 2A)**. Next, we tested other ferroptosis markers at different periods after SDDV infection. Compared to the control group, the intracellular levels of Fe^2+^ increased at 24 hpi and peaked at 48 hpi **(Fig. 2B)**, coinciding with excessive ROS accumulation **(Fig. 2C and D)**, leading to increased LPO levels **(Fig. 2E)**. In response, intracellular GSH was oxidized to glutathione disulfide (GSSG)to scavenge massive ROS under oxidative stress. Therefore, GSH and GPX4 was down-regulated in the middle and late stages of infection, reaching the lowest value at 60 hpi **(Fig. 2F and G)**. In conclusion, the above results showed that SDDV infection activates ferroptosis in MFF-1 cells.

**Fig. 2.**
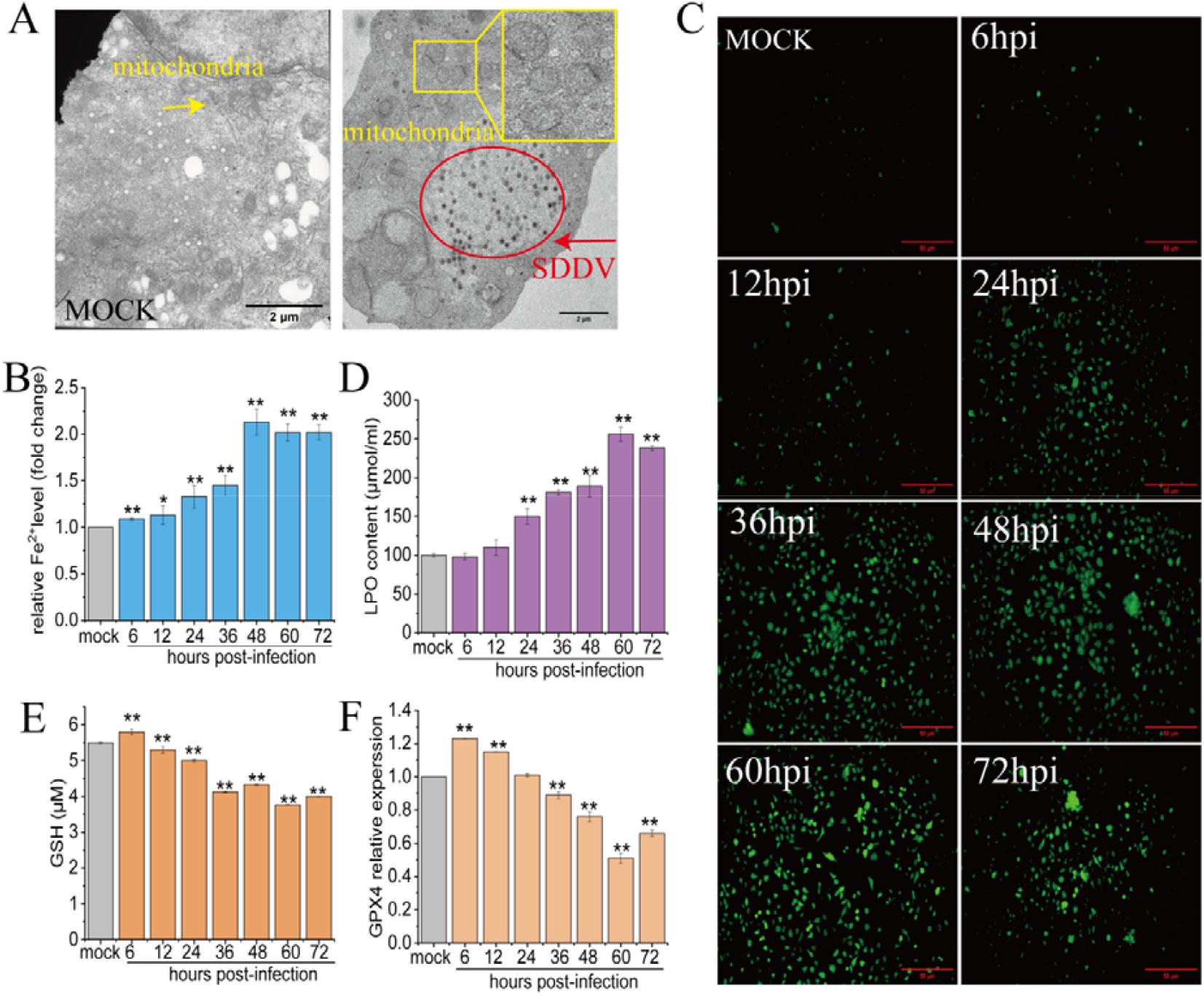
Ferroptosis contributes to SDDV-induced cell death in vitro. **A** Transmission electron microscopy images showing mitochondrial morphology and viral particles in control and SDDV-infected MFF-1 cells. Black arrows indicate mitochondria (scale bar = 1 μm). **B** Fe^2+^ concentrations were quantified using an iron assay kit, with absorbance measured at 593 nm. **C, D** ROS levels in MFF-1 cells were assessed through excitation wavelength of 488 nm and an emission wavelength of 525 nm and fluorescence intensity following SDDV infection. **E** LPO concentrations in MFF-1 cells were measured using an LPO assay kit. **F** GSH levels were quantified using a GSH assay kit, with absorbance measured at 412 nm. G The relative expression of GPX4 was evaluated in control and SDDV-infected MFF-1 cells by qRT-PCR. n = 3. *p < 0.05.

### 3.3. Ferroptosis is the primary form of cell death upon SDDV infection

Cell death caused by viral infection may involve a combination of different cell death pathways rather than a single mode [24]. Identifying the predominant cell death mode is essential. In this study, MFF-1 cells were treated with various concentrations of Fer-1 (a ferroptosis inhibitor), Erastin (a ferroptosis inducer), z-VAD (a pyroptosis inhibitor) and Nec1 (a necroptosis inhibitor) prior to SDDV infection, and the relative expression of the SDDV *mcp* gene was measured at 60 hpi. The results revealed that Fer-1 treatment exhibited the most significant inhibitory effect on SDDV replication (Fig. 3A), which was further confirmed by immunofluorescence assay (Fig. 3B). Treatment with Erastin produced the opposite effect (Fig. 3A and B). Notably, the treatments with z-VAD and Nec1 also significantly inhibited SDDV proliferation, suggesting that SDDV infection induces multiple forms of programmed cell death in MFF-1 cells. However, Fer-1 had the most pronounced inhibitory effect, indicating that ferroptosis is the dominant mode of cell death.

**Fig. 3.**
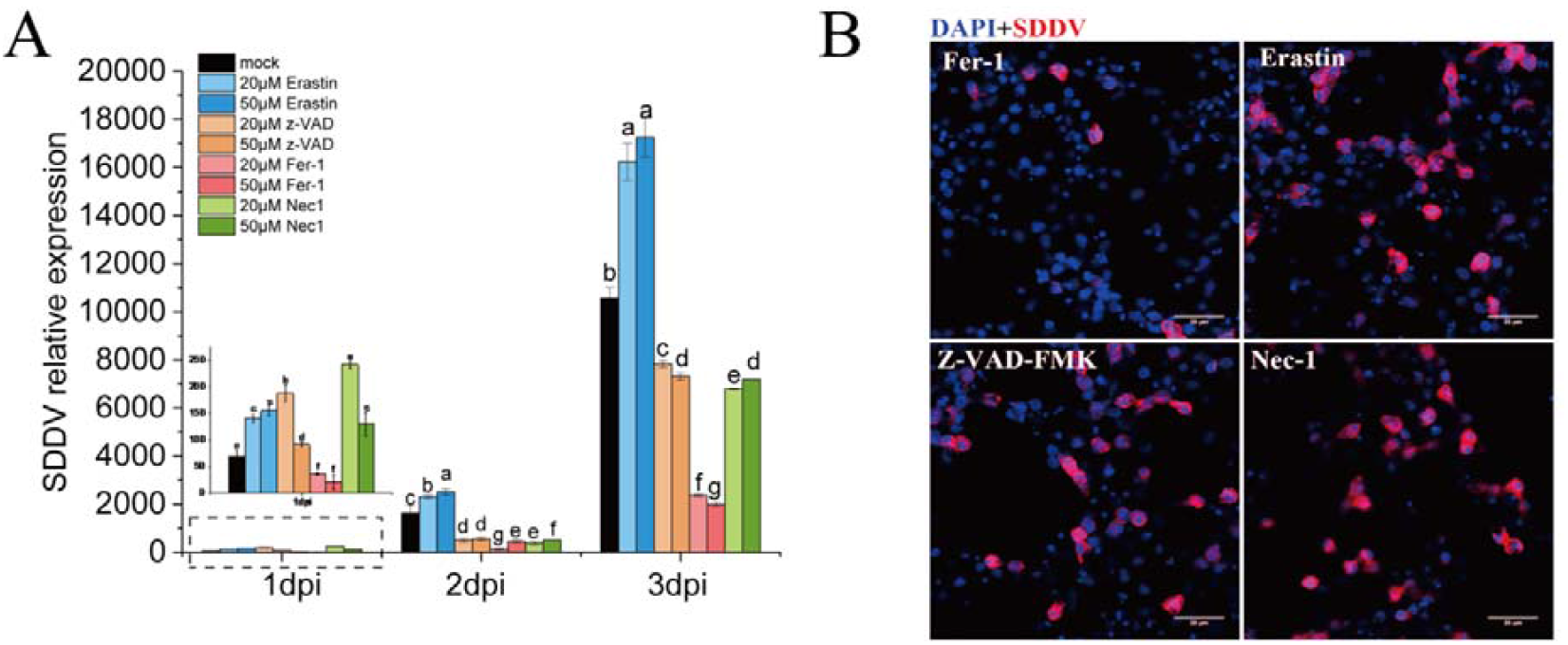
Ferroptosis is the primary mode of cell death induced by SDDV infection. **A** Relative SDDV expression levels in MFF-1 cells after treatment with 20 μM and 50 μM erastin, Fer-1, z-VAD, and Nec-1 for 4 h, followed by SDDV infection. The mock group was treated with diluted DMSO, Different letters (a, b, c, d) indicate significant differences between groups. **B** Immunofluorescence analysis of SDDV at 60 hpi in MFF-1 cells pre-treated with 50 μM Erastin, Fer-1, z-VAD, or Nec-1 for 5 h prior to SDDV challenge. n = 3.

### 3.4. Fer-1 treatment mitigated ferroptosis by SDDV infection

To examine whether the inhibition of ferroptosis can protect against SDDV-induced iron overload and lipid peroxidation, MFF-1 cells were treated with Fer-1, Erastin, or DMSO, followed by infection with SDDV at an MOI of 5.0. As shown in Figure 4A, DMSO and Erastin treatments induced accumulation of Fe^2+^, whereas Fer-1 treatment remarkably ameliorated such inductions during infection. We also assessed the levels of ROS and 8-OHdG in infected MFF-1 cells. After Fer-1 treatment, intracellular ROS **(Figures 4B and C)** and 8-OHdG **(Figure 4D)** content significantly declined compared to DMSO-treated cells, while Erastin treatment led to enhanced ROS and 8-OHdG production. Furthermore, GSH and GPX4 levels were reduced by Erastin treatment but restored following Fer-1 treatment **(Figures 4E and F)**. Importantly, we observed that SDDV-infected mandarinfish treated with Fer-1 exhibited lower mortality compared to those treated with Erastin or DMSO **(Figure 4G)**. These results demonstrate that Fer-1 can suppress ferroptosis induced by SDDV infection and reduce mortality in mandarinfish.

**Fig. 4.**
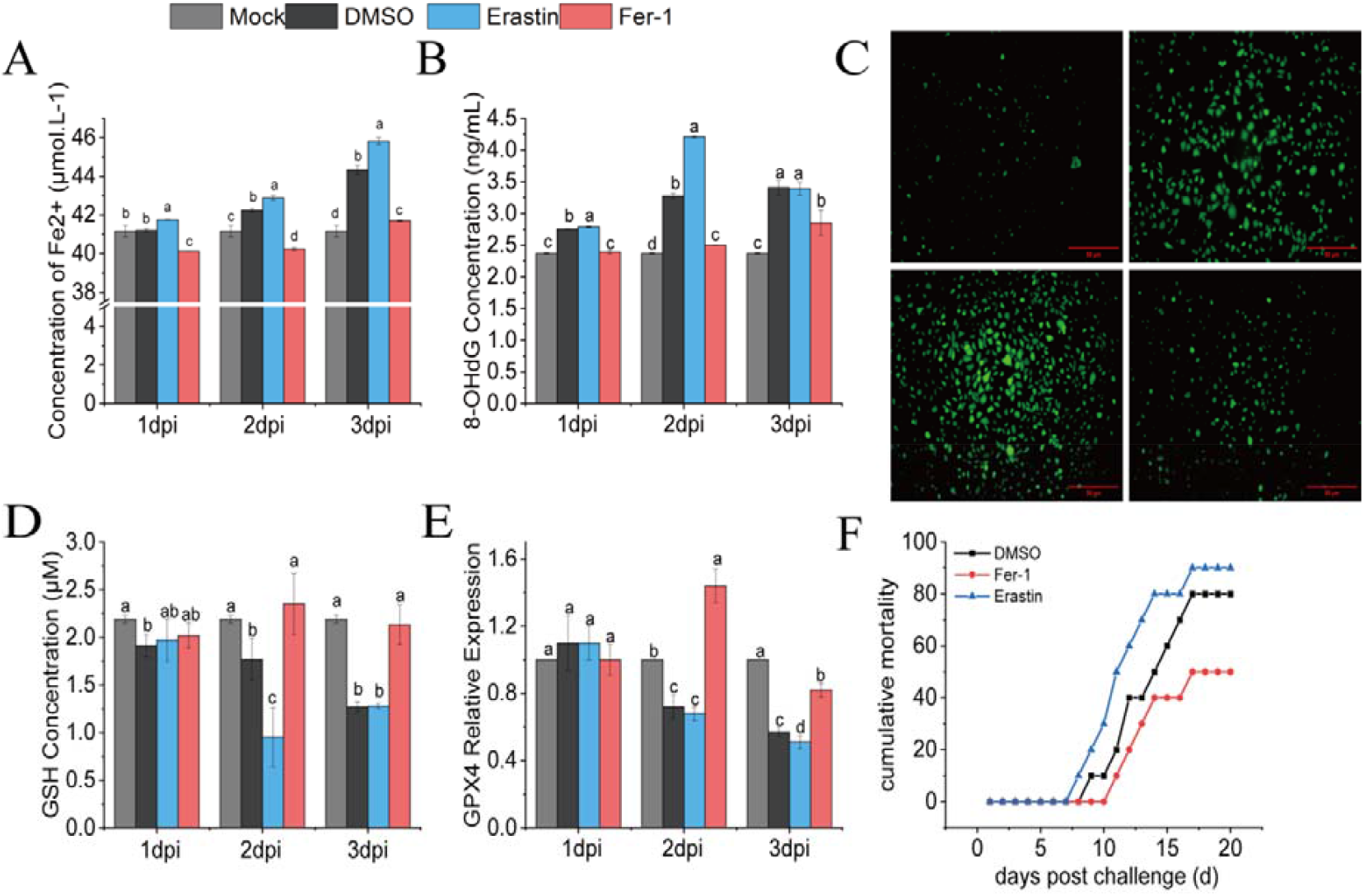
Fer-1 treatment alleviated ferroptosis induced by iron overload and lipid peroxidation following SDDV infection. Changes in Fe^2+^ concentration **(A)**, ROS levels (**B, C**), 8-OHdG content (**D**), and GSH content (**E**) in MFF-1 cells after SDDV infection. **F** The relative expression of GPX4 was analyzed by qRT-PCR in control and SDDV-infected MFF-1 cells. **G** Survival curve of mandarinfish after SDDV infection. Fish were intramuscularly injected with DMSO, erastin, or Fer-1, followed by SDDV challenge via intraperitoneal injection after 4 h. The mock group consisted of untreated mandarinfish, while the DMSO group was injected with diluted DMSO 5 h prior to SDDV infection, serving as a control for other drug-treated groups. *n* = 3. Different letters (a, b, c, d) indicate significant differences between groups.

### 3.5. SDDV-induced ferroptosis is iron dependent

Ferroptosis is driven by iron-dependent ROS production. To evaluate the effect of altering labile iron on SDDV-induced ferroptosis in vivo, mandarinfish were injected intramuscularly with DFO (an iron chelator, 0.1 g/kg), FAC (an iron supplement, 0.1 g/kg) or DMSO (control, PBS diluted to 20%) before infection with SDDV (10^3^ TCID^50^/fish). As a result, DFO treatment reduced the content of Fe3+ as observed in iron-stained spleen tissue sections (Fig. 5A). Moreover, ROS levels were significantly elevated in the FAC group compared to the DMSO control group in MFF-1 cells, while the DFO treatment group markedly reduced these levels (Fig. 5B). In the in vivo experiments, the LPO content was higher in the FAC group than in the control group, whereas the DFO group effectively reversed this increase (Fig. 5C). DFO treatment also restored GPx4 expression levels (Fig. 5D). Interestingly, DFO treatment protected mandarin fish from SDDV infection. As shown in Figure 5F, FAC treatment significantly increased the transcription of the SDDV *mcp* gene, while DFO treatment had the opposite effect. More importantly, mortality in the control and FAC-treated groups was 90% and 100%, respectively, whereas DFO treatment reduced mortality to 50% **(Fig. 5E)**. All together, these findings suggest that SDDV infection induces iron overload, ROS accumulation, and lipid peroxidation through an iron-dependent mechanism, and reducing iron levels improves the survival rate in mandarinfish infected with SDDV.

**Fig. 5.**
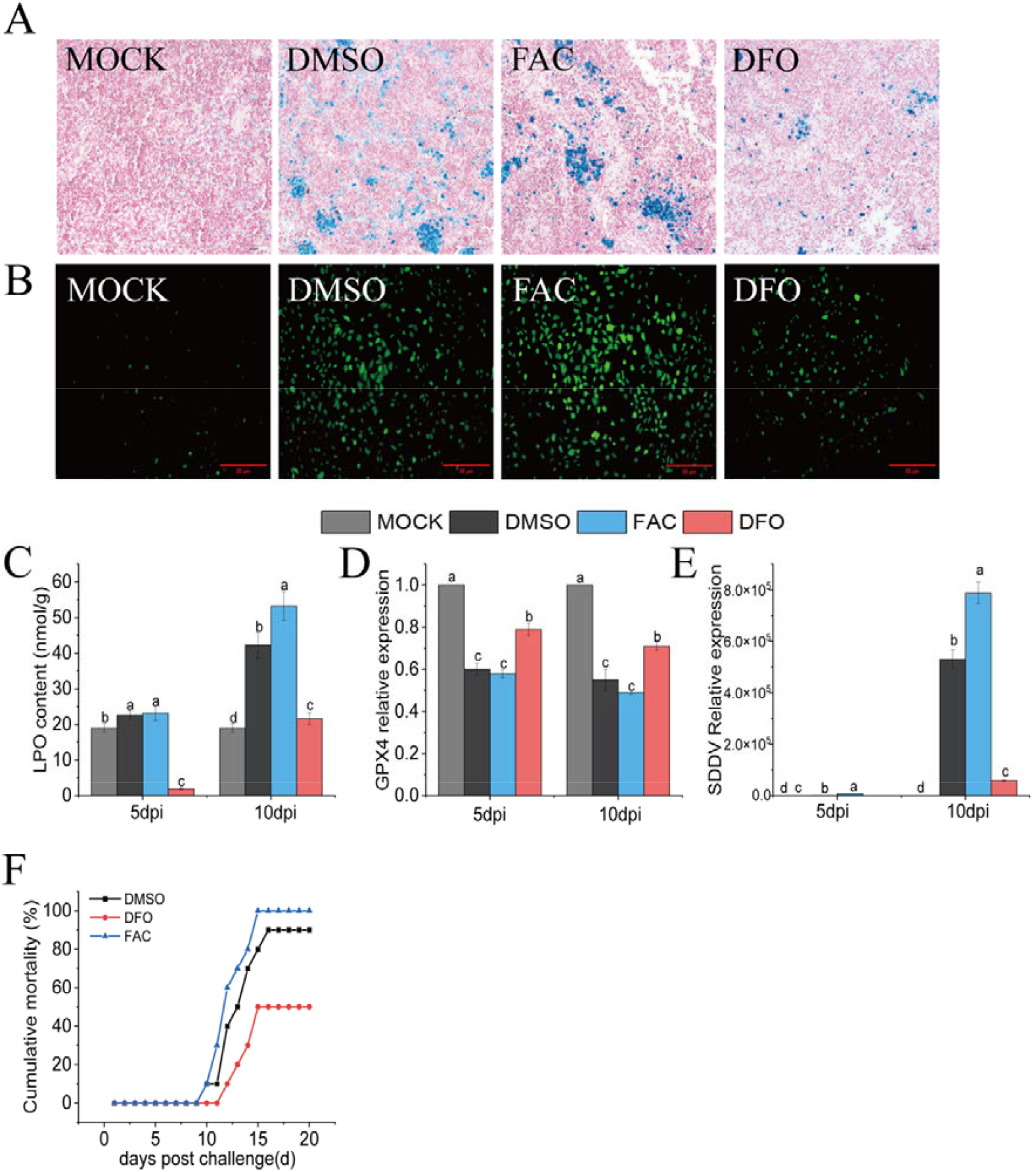
Regulating labile iron level in vivo affect SDDV infection. **A**, Representative images of Prussian blue staining in spleen tissues of mandarinfish from the mock, control, DFO, and FAC groups (scale bar = 50 μm). **B, C** Statistical analysis of lipid ROS and LPO levels showed a significant decrease in the DFO and FAC groups compared to the mock and DMSO group. **D** Relative mRNA expression of GPX4 was measured by qPCR in the DFO and FAC groups compared to the mock and DMSO group. **E** Mortality rates of mandarinfish were recorded after intraperitoneal injection with FAC and DFO, followed by SDDV challenge. **F** SDDV mRNA levels were quantified by qPCR after mandarinfish were injected with FAC and DFO and challenged with SDDV. The mock group consisted of untreated mandarinfish, while the DMSO group was injected with diluted DMSO 5 h prior to SDDV infection, serving as a control for other drug-treated groups. *n* = 3. The letters a, b, c, and d represent groups that have significant differences between them (*p* < 0.05).

### 3.6. TfR1 is involved in ferroptosis in SDDV-infected MFF-1 cells

The regulation of iron homeostasis affects cellular sensitivity to ferroptosis. To further explore the mechanisms underlying SDDV-induced ferroptosis, we focused on the genes for maintaining iron homeostasis, including TfR1, iron responsive element binding protein 2 (IREB2), nuclear receptor coactivator 4 (NCOA4) and ferritin Heavy Chain 1 (FTH1). IREB2 regulates cellular iron levels by modulating the translation and stability of mRNAs that control iron metabolism during iron depletion. FTH1 is a major component of ferritin, an iron storage protein [30]. NCOA4 mediates ferritin autophagy and the release of free iron, which is essential for inducing ferroptosis [31]. In the study, the results showed that SDDV infection significantly altered the expression of these genes in MFF-1 cells. In detail, the expression levels of NCOA4, TfR1 and IREB2 was significantly upregulated **(Fig. 6A, C and D)**, while FTH1 mRNA levels decreased at later stages of SDDV infection **(Fig. 6B)**. Among the genes analyzed, the most pronounced change was observed in mandarinfishTfR1 mRNA levels, which were significantly upregulated by more than fourfold at 60 hpi **(Fig. 6D)**. Consistently, Western blot analysis demonstrated a time-dependent increase in TfR1 protein levels in response to SDDV infection **(Fig. 6E)**. Moreover, our previous study identified that TfR1 is packaged into the SDDV mature virion through LC-MS/MS (Unpublished data). Based on this, we hypothesized that TfR1 might serve as a key effector gene in SDDV-induced ferroptosis. To test this hypothesis, we evaluated the effect of TfR1 knockdown. MFF-1 cells were transfected with either an irrelevant control siRNA or TfR1-specific siRNA. A significant decrease in both TfR1 mRNA and protein levels was observed in TfR1 knockdown cells compared with scrambled siRNA controls **(Fig. 6F, G**, and **H)**. Twenty-four hours post transfection, MFF-1 cells were infected with SDDV at an MOI of 5.0 and features of ferroptosis were assessed by monitoring the intracellular levels of Fe^2+^, ROS and LPO. As expected, Fe^2+^ level was lower in TfR1 knockdown cells compared to siRNA control cells (**Fig. 6I**). Correspondingly, the accumulation of LPO and ROS induced by SDDV infection was reduced in the TfR1 knockdown cells **(Fig. 6J** and **K)**. Additionally, we found that siTfR1 treatment enhanced cell viability during the SDDV infection process **(Fig. 6L)**, as measured by the CCK-8 assay. To further evaluate the direct impact of TfR1 knockdown on SDDV replication, the transcription level of the SDDV *mcp* gene was measured at 24 and 48 hpi via RT-qPCR. As shown in **Fig. 6M**, SDDV replication was suppressed following siTfR1 treatment. In conclusion, TfR1 plays a critical role in ferroptosis induced by SDDV infection, and downregulation of TfR1 inhibits ferroptosis in SDDV-infected MFF-1 cells.

**Fig. 6.**
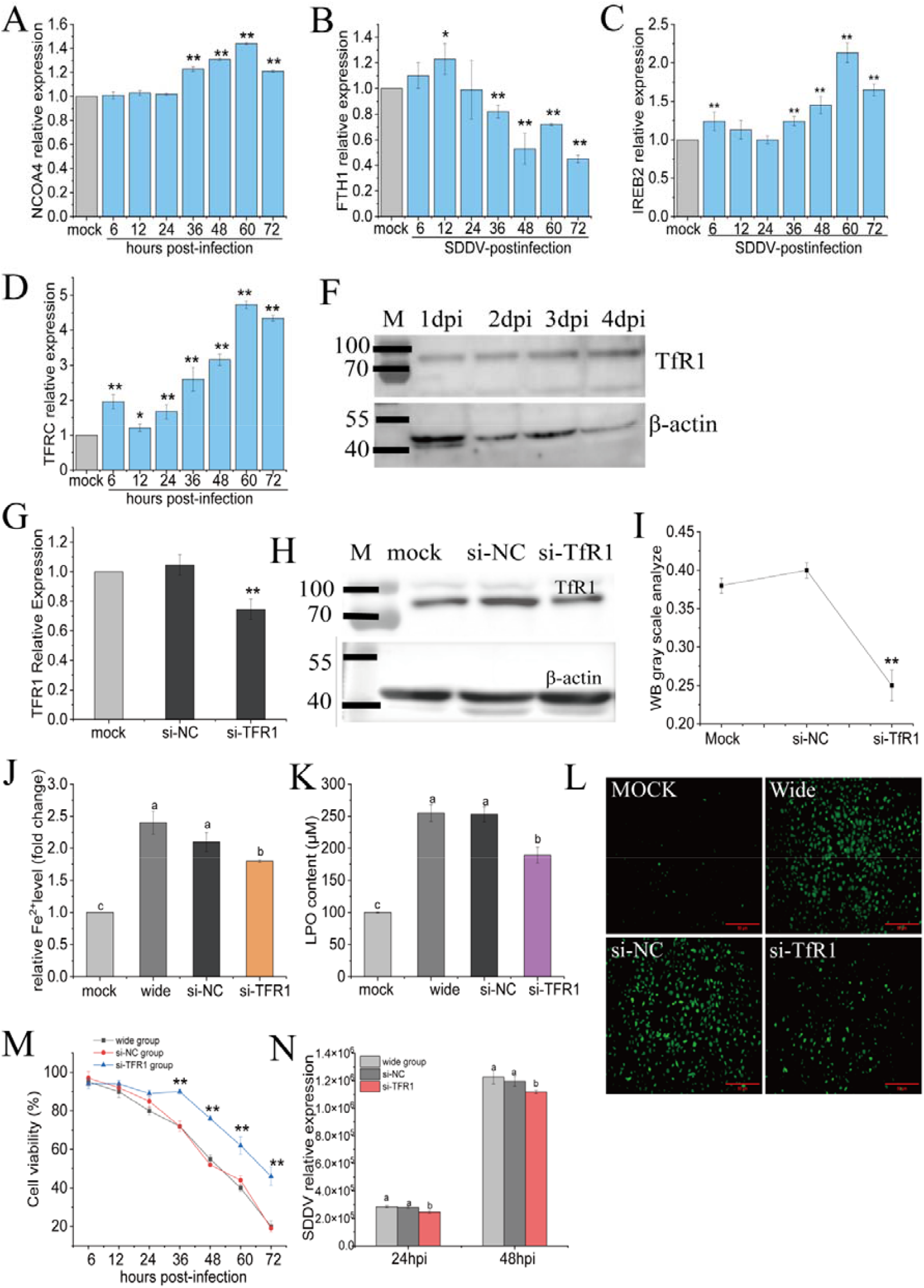
Downregulation of TFRC inhibited ferroptosis in SDDV-infected MFF-1 cells. **A-D** The mRNA levels of ferroptosis-related genes were detected by qPCR. All data are presented as mean ± SD. *p < 0.05, **p < 0.001. E The protein level of TfR1 was measured by western blotting. **F-H** The effects of TfR1 knockdown at both the mRNA level (via qPCR) and protein level (via western blotting). **I-K** Statistical analysis showed that Fe^2+^, LPO, and ROS levels significantly decreased in the siTfR1 group compared to the si-NC group (\**p* < 0.05, \*\**p* < 0.001). **L** Cell viability in the siTfR1 group was notably improved following SDDV infection, as determined by CCK-8 assays. **M** The relative expression of SDDV mRNA in the siTfR1 group was significantly lower than that in the si-NC group. The mock group refers to untreated cells, while the wide group represents cells infected with SDDV without any other treatment. *n* = 3. The letters a, b, c, and d indicate groups with statistically significant differences.

## 4. Discussion

Ferroptosis represents a newly recognized form of regulated cell death, primarily driven by iron-dependent lipid peroxidation. Although ferroptosis was only recently identified, increasing evidence highlights a strong association between viral infections and this unique cell death pathway. Ferroptosis is characterized by elevated iron levels, which promote the production of ROS via the Fenton reaction, thereby accelerating LPO [32, 33]. This study uncovers the potential role and underlying mechanism of ferroptosis during SDDV infection, providing new insights into the interaction between SDDV and its host.

Following SDDV infection, tannin staining of spleen tissue from mandarinfish revealed a phenomenon of free iron accumulation **(Fig. 1A)**. Additionally, the levels of LPO, ROS, and 8-OHdG in cells and tissues escalated, indicating the existence of oxidative stress after infection **(Fig. 1C and Fig. 2C, D, E)**. GPx4, a key intracellular antioxidant enzyme, plays a crucial role in regulating ferroptosis and its expression is suppressed during the intrinsic pathway of ferroptosis [23, 29]. GPx4 mitigates lipid peroxidation by utilizing GSH, thereby protecting cells from ferroptosis. In our study, we examined the alterations in the GSH cycle following SDDV infection and observed a downregulation of GPX4 expression, accompanied by a significant reduction in GSH levels. Furthermore, TEM images of SDDV-infected cells revealed mitochondria exhibiting typical morphological features of ferroptosis **(Fig. 2A)**. Fer-1, a potent inhibitor of ferroptosis, was shown to prevent the accumulation of LPO resulting from iron overload. In this study, Fer-1 treatment alleviated iron overload, reduced LPO and ROS accumulation, and prevented the decline in GSH and GPx4 in SDDV-infected MFF-1 cells. Additionally, Fer-1 treatment significantly decreased the mortality of mandarinfish infected with SDDV. In contrast, treatment with Erastin, a ferroptosis inducer, produced the opposite effect. These findings confirm that the observed changes are characteristic of ferroptosis, indicating its activation during SDDV infection.

Another two fish iridoviruses, ISKNV and MRV, were introduced for a comparative study. Upon infected with ISKNV, the content of LPO markedly reduced at 3 dpi **(Fig.1C)**. After MRV infection, 8-OHdG levels decreased and GSH levels increased. These results are inconsistent with the hallmark characteristics of ferroptosis, making it challenging to conclusively determine whether ferroptosis occurs after ISKNV or MRV infection. These observations suggest that the pathogenicity of ISKNV and MRV differs from that of SDDV.

Viral infection-induced cell death may involve a mixed pattern of cell death, rather than a single mode [24]. Our study demonstrated that SDDV infection in MFF-1 cells was suppressed by inhibitors of ferroptosis, apoptosis, and necrosis, suggesting that SDDV triggers multiple forms of cell death. However, the most pronounced inhibitory effect was observed with Fer-1 treatment, indicating that ferroptosis is the predominant form of cell death induced by SDDV infection.

Iron is an essential element required to support basic cellular processes, and it also plays a crucial role in the growth, virulence and pathogenicity of viruses, which acquire iron from the host. Notably, DFO, an iron-chelating agent that reduces free iron levels, has been identified as an effective modulator of ferroptosis and is well-documented for its antiviral activity [34]. Kannan et al. reported that pretreatment of mouse primary microglial cells with DFO blocked HIV-1 Tat-mediated microglial activation in vitro and reduced the expression and release of proinflammatory cytokines [35].Additionally, DFO has been shown to inhibit SARS-CoV-2 infection and block SARS-CoV-2-induced ferroptosis [36]. In our study, DFO administration to SDDV-infected mandarinfish led to a decrease in iron content, along with reductions in LPO and 8-OHdG levels, and increases in GPx4 and GSH. Moreover, DFO treatment significantly inhibited SDDV replication, resulting in a 50% reduction in mortality. These findings suggest that DFO prevents SDDV-induced ferroptosis and reduces viral replication by lowering iron content in vivo.[1]

Transferrin receptor 1 (TfR1) mediates cellular iron uptake and is essential for maintaining iron homeostasis [37]. Our results demonstrated that TfR1 expression was significantly increased in SDDV-infected MFF-1 cells at 72 hpi. Knockdown of TfR1 alleviated SDDV-induced ferroptosis in MFF-1 cells, leading to reduced iron overload, decreased LPO levels, upregulated GPX4 expression, and decreased viral replication. TfR1 has also been shown to play a vital role in the proliferation of several viruses. For example, TfR1 cooperates with metabotropic glutamate receptor subtype 2 to mediate the internalization of Rabies virus (RABV) and SARS-CoV-2 [38]; Porcine epidemic diarrhea virus (PEDV) S1 protein interacts with TfR1 to promote viral entry [39]. Several other viruses have been shown to exploit cellular TfR1 as a receptor, facilitating viral internalization through clathrin-mediated endocytosis, including New World hemorrhagic fever arenaviruses [40], Transmissible gastroenteritis virus (TGEV) [41] and mouse mammary tumor virus (MMTV) [42]. Furthermore, our previous data showed that high abundance of mandarinfish TfR1 existed in the purified SDDV virus particles. Therefore, we hypothesize that TfR1 may plays an important role in the process of SDDV infection. Future studies will further investigate how TfR1 regulates SDDV infection and which stages it influences.

## 5. Conclusion

In summary, SDDV infection induces ferroptosis both *in vivo* and *in vitro*. Ferroptosis, a form of iron-dependent cell death, is the predominant mode of cell death during SDDV infection. Mechanistically, the key iron-regulating gene TfR1 is upregulated by SDDV infection, leading to the release of free iron, which triggers lipid peroxidation and ferroptosis. Both the downregulation of TfR1 expression and the reduction of iron levels inhibit ferroptosis, thereby decreasing viral replication and lowering mortality in mandarinfish.

## Acknowledgments

This work was funded by key areas R&D Program of Guangdong Province under No. 2021B0202040002; Guangdong Basic and Applied Basic Research Foundation under No. 2023A1515110938; National Natural Science Foundation of China under No. 32403075; China Postdoctoral Science Foundation under No. 2024M753721.

## References

1. de Groof A, Guelen L, Deijs M, van der Wal Y, Miyata M, Ng KS, et al. A Novel Virus Causes Scale Drop Disease in Lates calcarifer. PLoS Pathog. 2015;11(8):e1005074. Epub 2015/08/08. doi: 10.1371/journal.ppat.1005074. PubMed PMID: 26252390; PubMed Central PMCID: PMCPMC4529248.

2. Kayansamruaj P, Soontara C, Dong HT, Phiwsaiya K, Senapin S. Draft genome sequence of scale drop disease virus (SDDV) retrieved from metagenomic investigation of infected barramundi, Lates calcarifer (Bloch, 1790). J Fish Dis. 2020;43(10):1287–98. Epub 2020/08/24. doi: 10.1111/jfd.13240. PubMed PMID: 32829517.

3. Nurliyana M, Lukman B, Ina-Salwany MY, Zamri-Saad M, Annas S, Dong HT, et al. First evidence of scale drop disease virus in farmed Asian seabass (Lates Calcarifer) in Malaysia) in Malaysia. Aquaculture. 2020;528. oi: ARTN 735600 10.1016/j.aquaculture.2020.735600. PubMed PMID: WOS:000553684000009.

4. Senapin S, Dong HT, Meemetta W, Gangnonngiw W, Sangsuriya P, Vanichviriyakit R, et al. Mortality from scale drop disease in farmed Lates calcarifer in Southeast Asia. J Fish Dis. 2019;42(1):119–27. Epub 2018/11/07. doi: 10.1111/jfd.12915. PubMed PMID: 30397913.

5. Gibson-Kueh S, Chee D, Chen J, Wang YH, Tay S, Leong LN, et al. The pathology of ‘scale drop syndrome’ in Asian seabass, Lates calcarifer Bloch, a first description. J Fish Dis. 2012;35(1):19–27. Epub 2011/11/23. doi: 10.1111/j.1365-2761.2011.01319.x. PubMed PMID: 22103767.

6. Fu Y, Li Y, Fu W, Su H, Zhang L, Huang C, et al. Scale Drop Disease Virus Associated Yellowfin Seabream (Acanthopagrus latus) Ascites Diseases, Zhuhai, Guangdong, Southern China: The First Description. Viruses. 2021;13(8). Epub 2021/08/29. doi: 10.3390/v13081617. PubMed PMID: 34452481; PubMed Central PMCID: PMCPMC8402775.

7. Fu W, Li Y, Fu Y, Zhang W, Luo P, Sun Q, et al. The Inactivated ISKNV-I Vaccine Confers Highly Effective Cross-Protection against Epidemic RSIV-I and RSIV-II from Cultured Spotted Sea Bass Lateolabrax maculatus. Microbiol Spectr. 2023;11(3):e0449522. Epub 2023/05/24. doi: 10.1128/spectrum.04495-22. PubMed PMID: 37222626; PubMed Central PMCID: PMCPMC10269448.

8. Dong C, Weng S, Luo Y, Huang M, Ai H, Yin Z, et al. A new marine megalocytivirus from spotted knifejaw, Oplegnathus punctatus, and its pathogenicity to freshwater mandarinfish, Siniperca chuatsi. Virus Res. 2010;147(1):98–106. Epub 2009/11/10. doi: 10.1016/j.virusres.2009.10.016. PubMed PMID: 19895861.

9. Fu YT, Li Y, Chen JM, Yu FZ, Liu XR, Fu WX, et al. A mandarinfish Siniperca chuatsi infection and vaccination model for SDDV and efficacy evaluation of the formalin-killed cell vaccine in yellowfin seabream Acanthopagrus latus. Aquaculture. 2023;570. oi: ARTN 739428 10.1016/j.aquaculture.2023.739428. PubMed PMID: WOS:000952139600001.

10. Halaly MA, Subramaniam K, Koda SA, Popov VL, Stone D, Way K, et al. Characterization of a Novel Megalocytivirus Isolated from European Chub (Squalius cephalus). Viruses. 2019;11(5). Epub 2019/05/18. doi: 10.3390/v11050440. PubMed PMID: 31096590; PubMed Central PMCID: PMCPMC6563503.

11. Shahin K, Subramaniam K, Camus AC, Yazdi Z, Yun S, Koda SA, et al. Isolation, Identification and Characterization of a Novel Megalocytivirus from Cultured Tilapia (Oreochromis spp.) from Southern California, USA. Animals (Basel). 2021;11(12). Epub 2021/12/25. doi: 10.3390/ani11123524. PubMed PMID: 34944299; PubMed Central PMCID: PMCPMC8697977.

12. Chinchar VG, Essbayer S, He JG, Hyatt A, Miyazaki T, Seligy V, et al. Family Iridoviridae. In: Fauquet CM, Mayo MA, Maniloff J, Desselberger U, Ball LA (eds) Virus taxonomy: 8th report of the International Committee on the Taxonomy of Viruses Elsevier Academic Press: San Diego, CA, USA. 2005:163–75.

13. Kurita J, Nakajima K. Review: Megalocytiviruses. Viruses. 2012;4:521–38.

14. Zhao L, Zhou X, Xie F, Zhang L, Yan H, Huang J, et al. Ferroptosis in cancer and cancer immunotherapy. Cancer Commun (Lond). 2022;42(2):88–116. Epub 2022/02/09. doi: 10.1002/cac2.12250. PubMed PMID: 35133083; PubMed Central PMCID: PMCPMC8822596.

15. Ursini F, Maiorino M. Lipid peroxidation and ferroptosis: The role of GSH and GPx4. Free Radic Biol Med. 2020;152:175–85. Epub 2020/03/14. doi: 10.1016/j.freeradbiomed.2020.02.027. PubMed PMID: 32165281.

16. Wang B, Shen WB, Yang P, Turan S. SARS-CoV-2 infection induces activation of ferroptosis in human placenta. Front Cell Dev Biol. 2022;10:1022747. Epub 2022/11/26. doi: 10.3389/fcell.2022.1022747. PubMed PMID: 36425527; PubMed Central PMCID: PMCPMC9679405.

17. Kan X, Yin Y, Song C, Tan L, Qiu X, Liao Y, et al. Newcastle-disease-virus-induced ferroptosis through nutrient deprivation and ferritinophagy in tumor cells. iScience. 2021;24(8):102837. Epub 2021/08/10. doi: 10.1016/j.isci.2021.102837. PubMed PMID: 34368653; PubMed Central PMCID: PMCPMC8326413.

18. Xu XQ, Xu T, Ji W, Wang C, Ren Y, Xiong X, et al. Herpes Simplex Virus 1-Induced Ferroptosis Contributes to Viral Encephalitis. mBio. 2023;14(1):e0237022. Epub 2022/12/13. doi: 10.1128/mbio.02370-22. PubMed PMID: 36507835; PubMed Central PMCID: PMCPMC9973258.

19. Ouyang A, Chen T, Feng Y, Zou J, Tu S, Jiang M, et al. The Hemagglutinin of Influenza A Virus Induces Ferroptosis to Facilitate Viral Replication. Adv Sci (Weinh). 2024;11(39):e2404365. Epub 2024/08/19. doi: 10.1002/advs.202404365. PubMed PMID: 39159143; PubMed Central PMCID: PMCPMC11497066.

20. Banerjee S, Sarkar R, Mukherjee A, Mitra S, Gope A, Chawla-Sarkar M. Rotavirus-induced lncRNA SLC7A11-AS1 promotes ferroptosis by targeting cystine/glutamate antiporter xCT (SLC7A11) to facilitate virus infection. Virus Res. 2024;339:199261. Epub 2023/11/06. doi: 10.1016/j.virusres.2023.199261. PubMed PMID: 37923170; PubMed Central PMCID: PMCPMC10684390.

21. Mao Q, Ma S, Li S, Zhang Y, Li S, Wang W, et al. PRRSV hijacks DDX3X protein and induces ferroptosis to facilitate viral replication. Vet Res. 2024;55(1):103. Epub 2024/08/19. doi: 10.1186/s13567-024-01358-y. PubMed PMID: 39155369; PubMed Central PMCID: PMCPMC11331664.

22. Zheng J, Conrad M. The Metabolic Underpinnings of Ferroptosis. Cell Metab. 2020;32(6):920–37. Epub 2020/11/21. doi: 10.1016/j.cmet.2020.10.011. PubMed PMID: 33217331.

23. Yi L, Hu Y, Wu Z, Li Y, Kong M, Kang Z, et al. TFRC upregulation promotes ferroptosis in CVB3 infection via nucleus recruitment of Sp1. Cell death & disease. 2022;13(7):592. Epub 2022/07/14. doi: 10.1038/s41419-022-05027-w. PubMed PMID: 35821227; PubMed Central PMCID: PMCPMC9276735.

24. Cheng J, Tao J, Li B, Shi Y, Liu H. Swine influenza virus triggers ferroptosis in A549 cells to enhance virus replication. Virol J. 2022;19(1):104. Epub 2022/06/18. doi: 10.1186/s12985-022-01825-y. PubMed PMID: 35715835; PubMed Central PMCID: PMCPMC9205082.

25. Dong C, Weng S, Shi X, Xu X, Shi N, He J. Development of a mandarin fish Siniperca chuatsi fry cell line suitable for the study of infectious spleen and kidney necrosis virus (ISKNV). Virus Res. 2008;135(2):273–81. Epub 2008/05/20. doi: 10.1016/j.virusres.2008.04.004. PubMed PMID: 18485510.

26. Zhu ZM, Duan C, Li Y, Huang CL, Weng SP, He JG, et al. Pathogenicity and histopathology of infectious spleen and kidney necrosis virus genotype II (ISKNV-II) recovering from mass mortality of farmed Asian seabass, Lates calcarifer, in Southern China. Aquaculture. 2021;534. doi: ARTN 736326 10.1016/j.aquaculture.2020.736326. PubMed PMID: WOS:000614762200002.

27. Zhang W, Deng H, Fu Y, Fu W, Weng S, He J, et al. Production and characterization of monoclonal antibodies against mandarinfish ranavirus and first identification of pyloric caecum as the major target tissue. J Fish Dis. 2023;46(3):189–99. Epub 2022/11/29. doi: 10.1111/jfd.13733. PubMed PMID: 36441809.

28. Zhang W, Gong H, Sun Q, Fu Y, Wu X, Deng H, et al. Peripheral B Lymphocyte Serves as a Reservoir for the Persistently Covert Infection of Mandarin Fish Siniperca chuatsi Ranavirus. Viruses. 2024;16(12). Epub 2025/01/08. doi: 10.3390/v16121895. PubMed PMID: 39772201; PubMed Central PMCID: PMCPMC11680134.

29. Dixon SJ, Lemberg KM, Lamprecht MR, Skouta R, Zaitsev EM, Gleason CE, et al. Ferroptosis: an iron-dependent form of nonapoptotic cell death. Cell. 2012;149(5):1060–72. Epub 2012/05/29. doi: 10.1016/j.cell.2012.03.042. PubMed PMID: 22632970; PubMed Central PMCID: PMCPMC3367386.

30. Scaramuzzino L, Lucchino V, Scalise S, Lo Conte M, Zannino C, Sacco A, et al. Uncovering the Metabolic and Stress Responses of Human Embryonic Stem Cells to FTH1 Gene Silencing. Cells. 2021;10(9). Epub 2021/09/29. doi: 10.3390/cells10092431. PubMed PMID: 34572080; PubMed Central PMCID: PMCPMC8469604.

31. Li JY, Feng YH, Li YX, He PY, Zhou QY, Tian YP, et al. Ferritinophagy: A novel insight into the double-edged sword in ferritinophagy-ferroptosis axis and human diseases. Cell Prolif. 2024;57(7):e13621. Epub 2024/02/23. doi: 10.1111/cpr.13621. PubMed PMID: 38389491; PubMed Central PMCID: PMCPMC11216947.

32. Frisk P, Tallkvist J, Gadhasson IL, Blomberg J, Friman G, Ilback NG. Coxsackievirus B3 infection affects metal-binding/transporting proteins and trace elements in the pancreas in mice. Pancreas. 2007;35(3):e37–44. Epub 2007/09/27. doi: 10.1097/mpa.0b013e3180986e84. PubMed PMID: 17895834.

33. Qiu C, Zhang X, Huang B, Wang S, Zhou W, Li C, et al. Disulfiram, a Ferroptosis Inducer, Triggers Lysosomal Membrane Permeabilization by Up-Regulating ROS in Glioblastoma. Onco Targets Ther. 2020;13:10631–40. Epub 2020/10/30. doi: 10.2147/OTT.S272312. PubMed PMID: 33116640; PubMed Central PMCID: PMCPMC7585819.

34. Wang J, Zhu J, Ren S, Zhang Z, Niu K, Li H, et al. The role of ferroptosis in virus infections. Frontiers in microbiology. 2023;14:1279655. Epub 2023/12/11. doi: 10.3389/fmicb.2023.1279655. PubMed PMID: 38075884; PubMed Central PMCID: PMCPMC10706002.

35. Kannan M, Sil S, Oladapo A, Thangaraj A, Periyasamy P, Buch S. HIV-1 Tat-mediated microglial ferroptosis involves the miR-204-ACSL4 signaling axis. Redox Biol. 2023;62:102689. Epub 2023/04/07. doi: 10.1016/j.redox.2023.102689. PubMed PMID: 37023693; PubMed Central PMCID: PMCPMC10106521.

36. Han Y, Zhu J, Yang L, Nilsson-Payant BE, Hurtado R, Lacko LA, et al. SARS-CoV-2 Infection Induces Ferroptosis of Sinoatrial Node Pacemaker Cells. Circ Res. 2022;130(7):963–77. Epub 2022/03/09. doi: 10.1161/CIRCRESAHA.121.320518. PubMed PMID: 35255712; PubMed Central PMCID: PMCPMC8963443.

37. Gammella E, Buratti P, Cairo G, Recalcati S. The transferrin receptor: the cellular iron gate. Metallomics. 2017;9(10):1367–75. Epub 2017/07/04. doi: 10.1039/c7mt00143f. PubMed PMID: 28671201.

38. Wang X, Wen Z, Cao H, Luo J, Shuai L, Wang C, et al. Transferrin Receptor Protein 1 Cooperates with mGluR2 To Mediate the Internalization of Rabies Virus and SARS-CoV-2. J Virol. 2023;97(2):e0161122. Epub 2023/02/14. doi: 10.1128/jvi.01611-22. PubMed PMID: 36779763; PubMed Central PMCID: PMCPMC9972945.

39. Zhang S, Cao Y, Yang Q. Transferrin receptor 1 levels at the cell surface influence the susceptibility of newborn piglets to PEDV infection. PLoS Pathog. 2020;16(7):e1008682. Epub 2020/07/31. doi: 10.1371/journal.ppat.1008682. PubMed PMID: 32730327; PubMed Central PMCID: PMCPMC7419007.

40. Radoshitzky SR, Abraham J, Spiropoulou CF, Kuhn JH, Nguyen D, Li W, et al. Transferrin receptor 1 is a cellular receptor for New World haemorrhagic fever arenaviruses. Nature. 2007;446(7131):92–6. Epub 2007/02/09. doi: 10.1038/nature05539. PubMed PMID: 17287727; PubMed Central PMCID: PMCPMC3197705.

41. Zhang S, Hu W, Yuan L, Yang Q. Transferrin receptor 1 is a supplementary receptor that assists transmissible gastroenteritis virus entry into porcine intestinal epithelium. Cell Commun Signal. 2018;16(1):69. Epub 2018/10/22. doi: 10.1186/s12964-018-0283-5. PubMed PMID: 30342530; PubMed Central PMCID: PMCPMC6196004.

42. Ross SR, Schofield JJ, Farr CJ, Bucan M. Mouse transferrin receptor 1 is the cell entry receptor for mouse mammary tumor virus. Proc Natl Acad Sci U S A. 2002;99(19):12386–90. Epub 2002/09/10. doi: 10.1073/pnas.192360099. PubMed PMID: 12218182; PubMed Central PMCID: PMCPMC129454.

